# Poultry farmer response to disease outbreaks in smallholder farming systems

**DOI:** 10.1101/2020.05.19.104059

**Authors:** Alexis Delabouglise, Nguyen Thi Le Thanh, Huynh Thi Ai Xuyen, Benjamin Nguyen-Van-Yen, Phung Ngoc Tuyet, Ha Minh Lam, Maciej F. Boni

**Affiliations:** Center for Infectious Diseases Dynamics, The Pennsylvania State University, University Park, PA, USA; Animals, Health, Territories, Risks, Ecosystems (ASTRE), Agricultural Research Centre for International Development (CIRAD), Montpellier, France; Oxford University Clinical Research Unit, Ho Chi Minh City, Vietnam; Ca Mau sub-Department of Livestock Production and Animal Health, Ca Mau, Vietnam; École Normale Supérieure, Paris, France; Centre for Tropical Medicine and Global Health, Nuffield Department of Medicince, University of Oxford, Oxford, United Kingdom

**Keywords:** epidemiology, poultry, avian influenza, Southeast Asia, behavioral epidemiology, health behavior, health economics, vaccination

## Abstract

Avian influenza outbreaks have been occurring on smallholder poultry farms in Asia for two decades. Farmer responses to these outbreaks can slow down or accelerate virus transmission. We used a longitudinal survey of 53 small-scale chicken farms in southern Vietnam to investigate the impact of outbreaks with disease-induced mortality on harvest rate, vaccination, and disinfection behaviors. We found that in small broiler flocks (≤16 birds/flock) the estimated probability of harvest was 56% higher when an outbreak occurred, and 214% higher if an outbreak with sudden deaths occurred in the same month. Vaccination and disinfection were strongly positively correlated with flock size and farm size, respectively. Small-scale farmers – the overwhelming majority of poultry producers in low-income countries – tend to rely on rapid sale of birds to mitigate losses from diseases. As depopulated birds are sent to markets or trading networks, this reactive behavior has the potential to enhance onward transmission.

**One sentence summary:** A cohort study of fifty three small-scale poultry farms in southern Vietnam reveals that when outbreaks occur with symptoms similar to highly pathogenic avian influenza, farmers respond by sending their chickens to market early, potentially exacerbating the effects of the outbreak.

## Introduction

Livestock production systems have been a major driver of novel pathogen emergence events over the past two decades (*1-3*). The conditions enabling the emergence and spread of a new disease in the human population partly depend on human behavioral changes, like hygiene improvements or social distancing, in the face of epidemiological risks (*4*). The same observation applies to disease emergence and spread in livestock populations as farmers adapt their farm management to maximize animal production and welfare while limiting cost in a constantly changing ecological and economic environment (*5*).

Poultry farming generates substantial risk for disease emergence. It is now the most important source of animal protein for the human population and the industry is changing rapidly (*6*). The link between poultry sector expansion and pathogen emergence is exemplified by the worldwide spread of the highly pathogenic form of avian influenza (AI) due to the H5N1 subtype of influenza A, after its initial emergence in China in 1996 (*2, 7*). Highly Pathogenic Avian Influenza (HPAI) causes severe symptoms in the most vulnerable bird species (including chicken, turkey, and quail), with mortality rates as high as 100% reported in broiler flocks (*8*). HPAI causes outbreaks in humans, and the risk that the pathogen makes the leap to a human pandemic is a persistent threat to public health (*9*). While HPAI does not persist in poultry populations in most affected countries, it has become endemic in parts of Asia and Africa and is periodically re-introduced into other areas like Europe and North America (*10, 11*). In affected countries, major factors influencing HPAI epidemiology appear to be farm disinfection, poultry vaccination, and marketing of potentially infected birds through trade networks, all of which depend on farmers’ management decisions (*12-16*).

It is still unclear how and to what extent changes in outbreak risk or mortality risk affect the behavior of poultry farmers. An anthropological study in Cambodia showed that high levels of farmer risk awareness associated with HPAI did not translate into major changes in their farming practices (*17*). Qualitative investigations conducted in Vietnam, Bangladesh, China, and Indonesia reported that farmers sometimes urgently sell or cull diseased poultry flocks as a way to mitigate economic losses, but evidence of this behavior’s onward epidemiological impact was not available (*12, 18-21*). Additionally, it is unknown whether poultry farmers increase application of disinfection practices or vaccination rates against avian influenza in response to disease outbreaks occurring in their flocks. Changes in farm management caused by variations in epidemiological risk have not been quantified for any livestock system that we are aware of, primarily because of the lack of combined epidemiological and behavioural data in longitudinal studies of livestock disease (*22*). Ifft et al. compared the evolution of chicken farm sizes and disease prevention in administrative areas with different levels of HPAI prevalence in Vietnam (*23*), and Hidano et al. modelled the effect of cattle mortality and production performance on the frequency of sales and culling in New Zealand dairy farms (*24*). One limitation of these two studies is that the dynamics were observed over long time steps (1 year), which does not allow for a precise estimation of the timing of farmer response after the occurrence of disease outbreaks and the potential feedback effect of this response onto the resulting outbreaks or epidemics.

We present a cohort study of small-scale poultry farms where we aimed to characterize the effect of disease outbreaks on livestock harvest rate and on two prevention practices, vaccination and farm disinfection. This longitudinal farm survey was conducted on small-scale poultry farms in the Mekong river delta region of southern Vietnam (*25*). HPAI has been endemic in this region since its initial emergence in 2003-2004 (*26*). Small-scale poultry farming is practiced by more than seven million Vietnamese households, mostly on a scale of fewer than 100 birds per farm (*27*). Farm production in this sector is severely impacted by infectious poultry diseases. Aside from HPAI, Newcastle disease, fowl cholera, and Gumboro are all endemic despite the availability of vaccines for their control (*28*).

## Materials and Methods

### 1. Data collection

An observational cohort study of small-scale poultry farms was conducted in Ca Mau province in southern Vietnam (*25, 29*). Fifty poultry farms from two rural communes were initially enrolled and three additional farms were subsequently added to the sample in order to replace three farmers who stopped their poultry farming activity. The two communes were chosen based on their high poultry density and their history of past HPAI outbreaks (*29*). Study duration was 20 months, from June 2015 to January 2017. Monthly Vietnamese-language questionnaires were used to collect information on (1) number of birds of each species and production type, (2) expected age of removal from the farm, (3) number of birds introduced, removed, and deceased in the last month, (4) clinical symptoms associated with death, (5) vaccines administered, (6) type of poultry housing used, and (7) disinfection activity. Each farm’s poultry were classified into “flocks”, defined as groups of birds of the same age, species, and production type (*25*). The main poultry species raised in these farms were chickens, ducks and Muscovy ducks.

### 2. Selection of data and covariates

We fit models with three different dependent variables: a “harvest model” of the probability of harvesting (i.e. selling or slaughtering) flocks at a particular production stage (data points are flock-months), an “AI vaccination model” of the probability of performing AI vaccination on flocks which had never received AI vaccination (data points are flock-months), and a “disinfection model” of the probability of disinfecting farm facilities (data points are farm-months). For the two first models, we focused our analysis on broiler chicken flocks. Chicken was the predominant species in the study population, the overwhelming majority of chicken flocks were broilers, and their age-specific harvest was easier to predict than the harvest of layer-breeder hens. Additionally, only six layer-breeder chicken flocks were vaccinated against AI during the study period. More details on data selection are provided in the **supplementary materials and methods 1**.

Outbreak categorical variables were included in each model, corresponding to an outbreak occurring in the corresponding farm in the same month, one month prior, and two months prior. An outbreak was defined as the death of at least two birds of the same species – on the same farm, in the same month, with similar clinical symptoms – as this may indicate the presence of an infectious pathogen on the farm. For the harvest model, only outbreaks in chickens were considered. For the AI vaccination model, outbreaks in chickens and outbreaks in any other species were included as two separate covariates. For the disinfection model, outbreaks in any of the species present in the farm were considered. In chickens, outbreaks with “sudden deaths” (i.e. the death of chickens in less than 24h after the onset of clinical symptoms) are considered as suspicions of highly pathogenic avian influenza (*30*). Therefore, we created two sub-categorical variables for outbreaks in chickens, with sudden deaths (OS, “outbreaks sudden”) and with no sudden deaths (ONS, “outbreaks not sudden”). For both the harvest and AI vaccination models, we assumed the effect of outbreaks on the dependent variable may be affected by the size of the considered flock (*n*). Consequently, we included this interaction term in the analysis.

The three dependent variables are likely affected by several farm-, flock-, and time-related factors, justifying the inclusion of several control covariates in the multivariate models. For the harvest model, we used the observed age of the chicken flocks (*t*) and the reported “intended age at harvest” (*t\**) and used *δt = t* – *t\** as the key independent variable indicating whether a flock was observed before or after its intended harvest time. For the AI vaccination model, the age of the chicken flocks *t* was considered as influencing the likelihood of vaccination. Other control covariates included flock size, age, calendar time (*T*), housing, vaccination status, introduction of other flocks onto the farm, and farm size (i.e. a farm’s poultry population size). The farm poultry population was broken down by species (chicken, duck, and Muscovy duck) and type of production (broiler and layer-breeder). Summary statistics of variables are displayed in the **Table 1** and all control covariates are listed in the **supplementary materials and methods 2**.

**Table 1.**
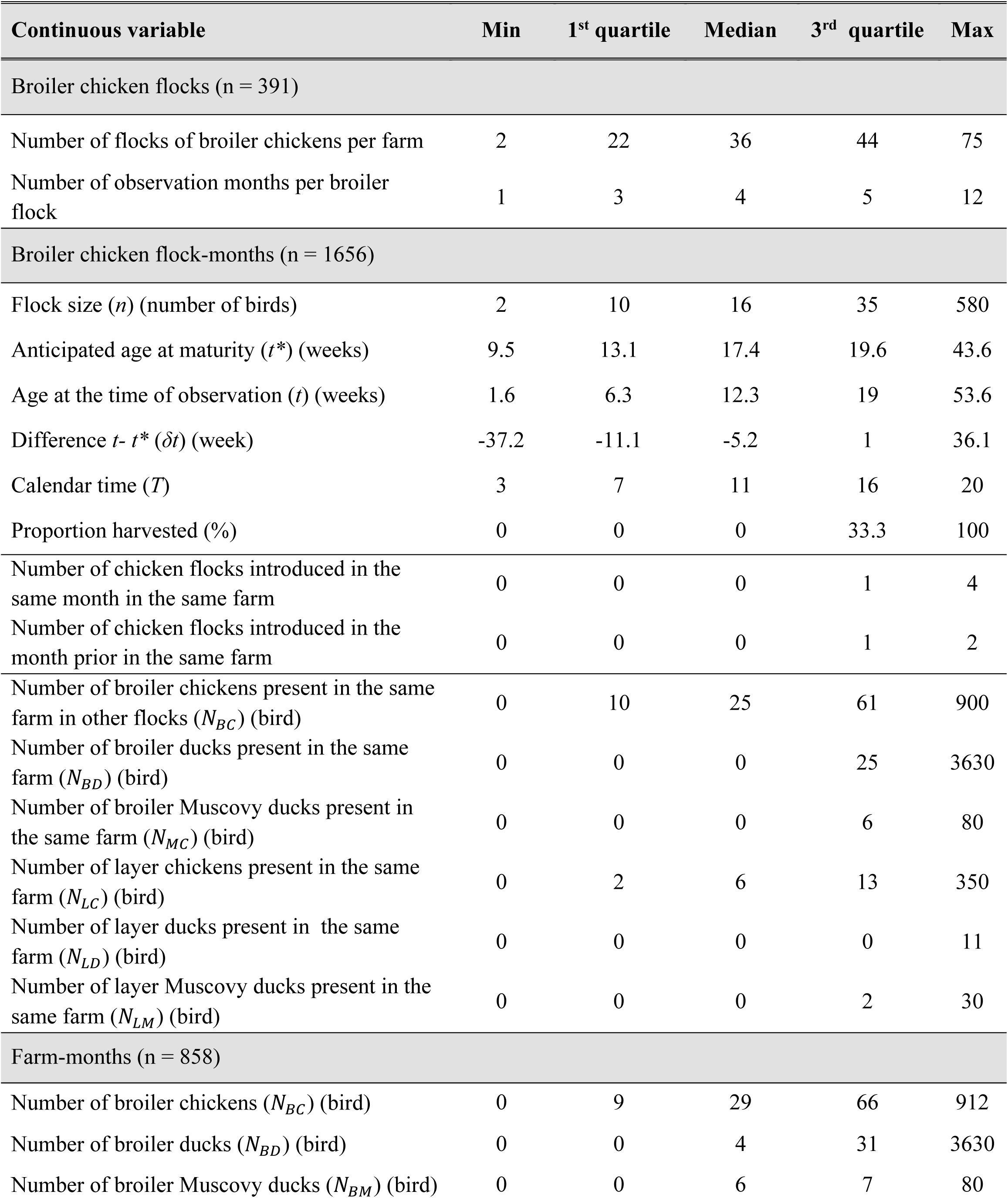
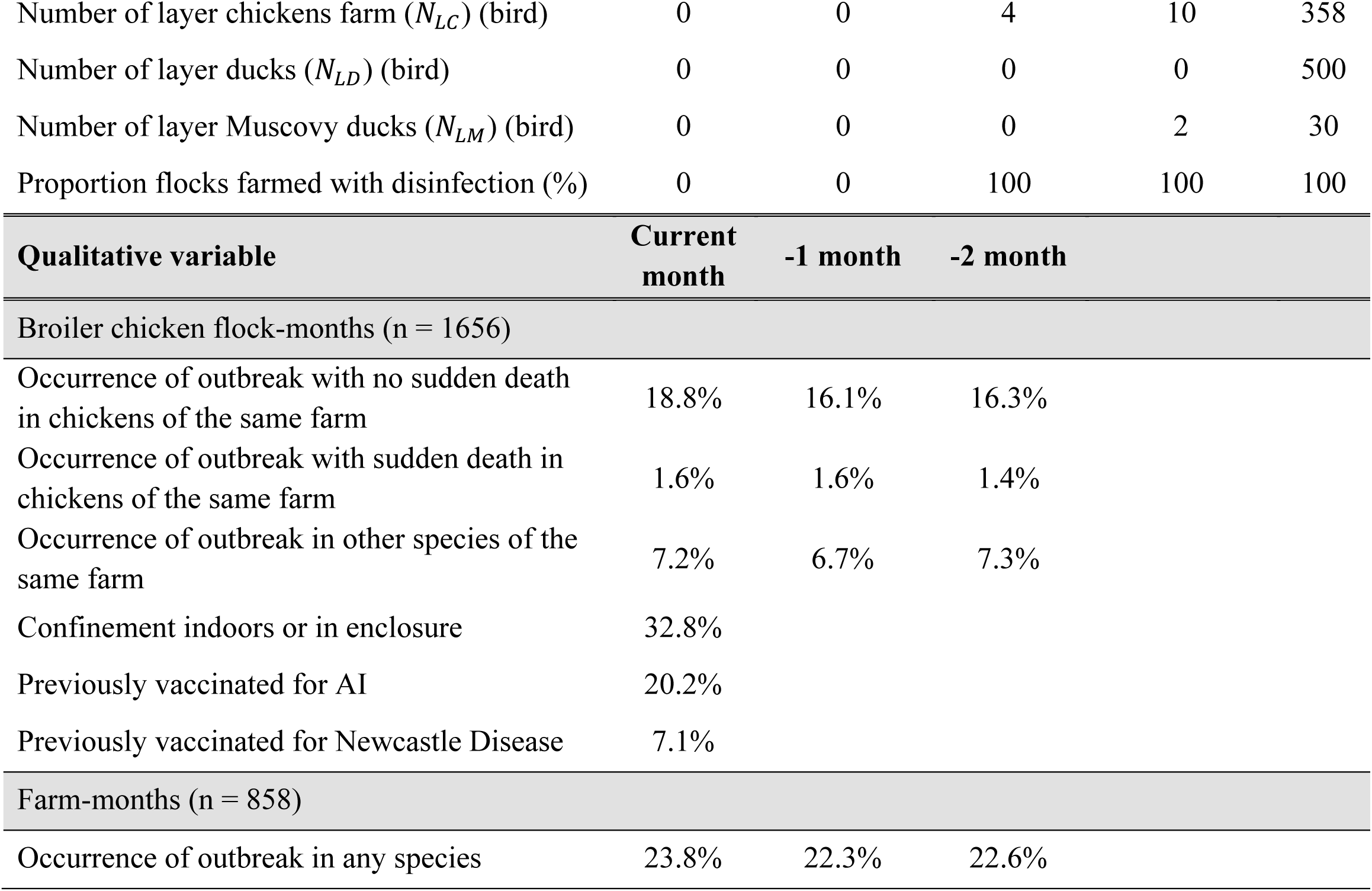
Summary statistics of variables.

| Continuous variable | Min | 1<sup>st</sup> quartile | Median | 3<sup>rd</sup> quartile | Max |
| --- | --- | --- | --- | --- | --- |
| Broiler chicken flocks (n = 391) |  |  |  |  |  |
| Number of flocks of broiler chickens per farm | 2 | 22 | 36 | 44 | 75 |
| Number of observation months per broiler flock | 1 | 3 | 4 | 5 | 12 |
| Broiler chicken flock-months (n = 1656) |  |  |  |  |  |
| Flock size ($n$) (number of birds) | 2 | 10 | 16 | 35 | 580 |
| Anticipated age at maturity ($t^*$) (weeks) | 9.5 | 13.1 | 17.4 | 19.6 | 43.6 |
| Age at the time of observation ($t$) (weeks) | 1.6 | 6.3 | 12.3 | 19 | 53.6 |
| Difference $t - t^*$ ($\delta t$) (week) | -37.2 | -11.1 | -5.2 | 1 | 36.1 |
| Calendar time ($T$) | 3 | 7 | 11 | 16 | 20 |
| Proportion harvested (%) | 0 | 0 | 0 | 33.3 | 100 |
| Number of chicken flocks introduced in the same month in the same farm | 0 | 0 | 0 | 1 | 4 |
| Number of chicken flocks introduced in the month prior in the same farm | 0 | 0 | 0 | 1 | 2 |
| Number of broiler chickens present in the same farm in other flocks ($N_{BC}$) (bird) | 0 | 10 | 25 | 61 | 900 |
| Number of broiler ducks present in the same farm ($N_{BD}$) (bird) | 0 | 0 | 0 | 25 | 3630 |
| Number of broiler Muscovy ducks present in the same farm ($N_{MC}$) (bird) | 0 | 0 | 0 | 6 | 80 |
| Number of layer chickens present in the same farm ($N_{LC}$) (bird) | 0 | 2 | 6 | 13 | 350 |
| Number of layer ducks present in the same farm ($N_{LD}$) (bird) | 0 | 0 | 0 | 0 | 11 |
| Number of layer Muscovy ducks present in the same farm ($N_{LM}$) (bird) | 0 | 0 | 0 | 2 | 30 |
| Farm-months (n = 858) |  |  |  |  |  |
| Number of broiler chickens ($N_{BC}$) (bird) | 0 | 9 | 29 | 66 | 912 |
| Number of broiler ducks ($N_{BD}$) (bird) | 0 | 0 | 4 | 31 | 3630 |
| Number of broiler Muscovy ducks ($N_{BM}$) (bird) | 0 | 0 | 6 | 7 | 80 |

| Number of layer chickens farm ($N_{LC}$) (bird) | 0 | 0 | 4 | 10 | 358 |
| --- | --- | --- | --- | --- | --- |
| Number of layer ducks ($N_{LD}$) (bird) | 0 | 0 | 0 | 0 | 500 |
| Number of layer Muscovy ducks ($N_{LM}$) (bird) | 0 | 0 | 0 | 2 | 30 |
| Proportion flocks farmed with disinfection (%) | 0 | 0 | 100 | 100 | 100 |
| Qualitative variable | Current month | -1 month | -2 month |  |  |
| Broiler chicken flock-months (n = 1656) |  |  |  |  |  |
| Occurrence of outbreak with no sudden death in chickens of the same farm | 18.8% | 16.1% | 16.3% |  |  |
| Occurrence of outbreak with sudden death in chickens of the same farm | 1.6% | 1.6% | 1.4% |  |  |
| Occurrence of outbreak in other species of the same farm | 7.2% | 6.7% | 7.3% |  |  |
| Confinement indoors or in enclosure | 32.8% |  |  |  |  |
| Previously vaccinated for AI | 20.2% |  |  |  |  |
| Previously vaccinated for Newcastle Disease | 7.1% |  |  |  |  |
| Farm-months (n = 858) |  |  |  |  |  |
| Occurrence of outbreak in any species | 23.8% | 22.3% | 22.6% |  |  |

### 3. Multivariate modelling

We assumed that the events of interest, namely harvest, AI vaccination, and disinfection were drawn from a binomial distribution and used a logistic function to link their probability to a function of the independent covariates. Some of the included effects are non-linear in nature, and we needed to account for the intra-farm autocorrelation of the dependent variables. We therefore used a mixed-effects general additive model (MGAM) implemented in R with the “mgcv” package (*31*). This enabled us to model the combined effect of *δt, t\**, and flock size (*n*) on harvest time; the effect of *t* and *n* on AI vaccination; and the effect of calendar time (*T*) on all the dependent variables, as penalized thin plate regression splines (*32*). We specifically chose these variables because they are presumably the most important factors influencing the dependent variables and their effect could possibly be highly non-linear. All other covariates were included as parametric regression terms. We also modelled the individual effects of farms on the dependent variables as random effects.

The complete models linking the logit *Y*_*ij*_ of probability of realization of an event and the set of explanatory variables, for a flock-month *i* (harvest, vaccination for AI) or a farm-month *i* (disinfection) in a farm *j*, are described by the following set of equations:

Harvest model (flock-month level):

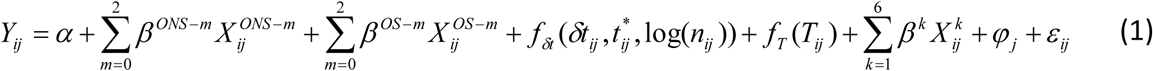

AI vaccination model (flock-month level):

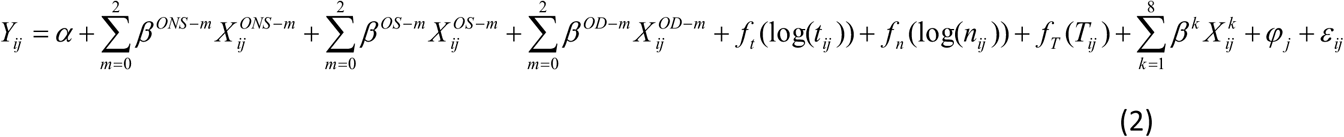

Disinfection model (farm-month level):

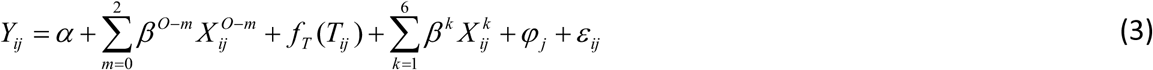

The model parameters are *α* the model intercept; *β* the parametric coefficients; *f* a thin-plate spline function; *X*^*k*^ the general notation for variables with linear effects; *X*^*O-m*^, *X*^*OS-m*^, *X*^*ONS-m*^ and *X*^*OD-m*^, categorical variables denoting presence or absence of an outbreak in the same farm *m* months prior in any species (O), in chickens with sudden deaths (OS), in chickens with no sudden deaths (ONS), and in different species (OD) respectively; *n* the flock size; *t* the current age of the flock; *t\** the age at maturity of the flock anticipated by the farmer; *δt* the difference between current age and age at maturity; *T* the calendar time; *φ* the farm random effect; *ε* the residual error term. Interaction terms between outbreak categorical variables and flock size *log(n*_*ij*_) were added in the Harvest and AI vaccination models.

Some variables with a highly skewed distribution were transformed (either log- or square-root-transformed). Details are provided in the **supplementary materials and methods 3**. Excessive multi-collinearity between covariates was assessed by estimating their variance inflated factor using the “usdm” R package (*33*). We fitted the complete models using the whole set of covariates using restricted maximum likelihood estimation. We then used a backward-forward stepwise selection, based on Akaike Information Criterion (AIC) comparison, to eliminate the variables with non-significant effects (*34*). Arguably, one farmer is likely to maintain the same farm management from one month to the next despite changes in influential covariates. Therefore, for each model, we tested the presence of farm-level time autocorrelation in the model deviance residuals. If there was sufficient evidence for the presence of autocorrelation, we implemented the same model fitting protocol with an additional intra-farm AR-1 time autocorrelation term on the dependent variable (*32*). More details on testing and accounting for time autocorrelation are provided in the **supplementary materials and methods 4.**

All analyses and graphical representations were performed with R version 3.6.1 (*35*).

### 4. Ethical statement

The collaboration between the investigators (authors) and the Ca Mau sub-department of livestock production and animal health (CM-LPAH) was approved by The Hospital for Tropical Diseases in Ho Chi Minh City, Vietnam. The CM-LPAH, which the equivalent of an ethics committee for studies dealing with livestock farming, specifically approved this study; at the province-level.

## Results

Fifty three farms were monitored from June 2015 to January 2017. **Figure 1** illustrates the farms’ structure and dynamics. Farmers kept an average number of 79 chickens, 53 ducks and 7 Muscovy ducks over the 20-month study period. Broiler chicken flocks were kept for 15.5 weeks on average after which most chickens were sold and a minority was consumed or kept on the farm for breeding and egg production.

**Figure 1.**
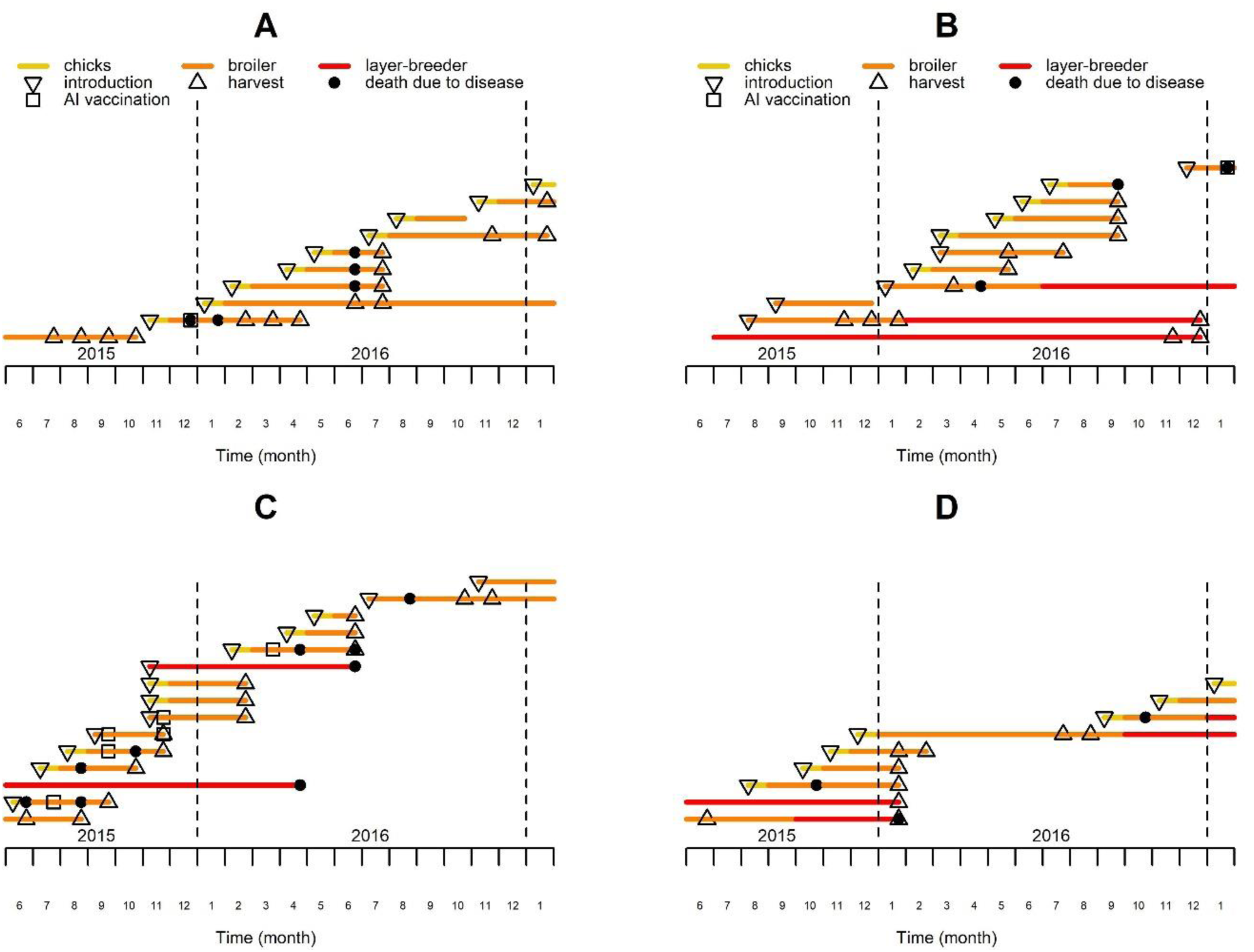
History of chicken flocks present in four of the observed farms over the study period. Each colored line represents the period over which a single chicken flock was present on the farm, with the color code indicating the production type, which may vary in the course of the flock production period. The major events affecting the flocks are located with specific symbols on the corresponding lines and months.

A total of 1656 broiler chicken flock-months were available for analysis. They belonged to 391 chicken flocks present on 48 farms. Occurrences of outbreaks in monitored farms in the same month, 1 month prior, and 2 months prior are summarized in **Table 1**. Additional descriptive statistics on control covariates are also displayed in **Table 1**. Out of this initial number, 1503 flock-months were selected for the harvest analysis (**Supplementary materials and methods 1**). No harvest occurred in 995 flock-months, complete harvest occurred in 258 flock-months, and partial harvest occurred in 250 flock-months. The probability of harvest during a month, with partial harvests weighted appropriately, was 23.9%. 1318 flock-months were selected for the AI vaccination analysis (**Supplementary materials and methods 1**). AI vaccination was performed in 7.5% (99/1318) of flock-months. The 99 vaccinated flocks were from 29 different farms (out of 48 farms keeping broiler chickens). For the disinfection model, 858 farm-months belonging to 52 farms were included (**Supplementary materials and methods 1**). During 552 farm-months the farm was fully disinfected, during 259 farm-months the farm was not disinfected at all, and during 47 farm-months disinfection was only performed for some of the flocks present in the farm. The probability of disinfection during a month, with partial disinfections weighted appropriately, was 67.4%. The best fit statistical models and their parameter values are summarized in **Table 2**. Fitted spline functions cannot be elegantly summarized by their coefficients and are most adequately represented with graphs (in **Figures 2 and 3**).

**Table 2.**
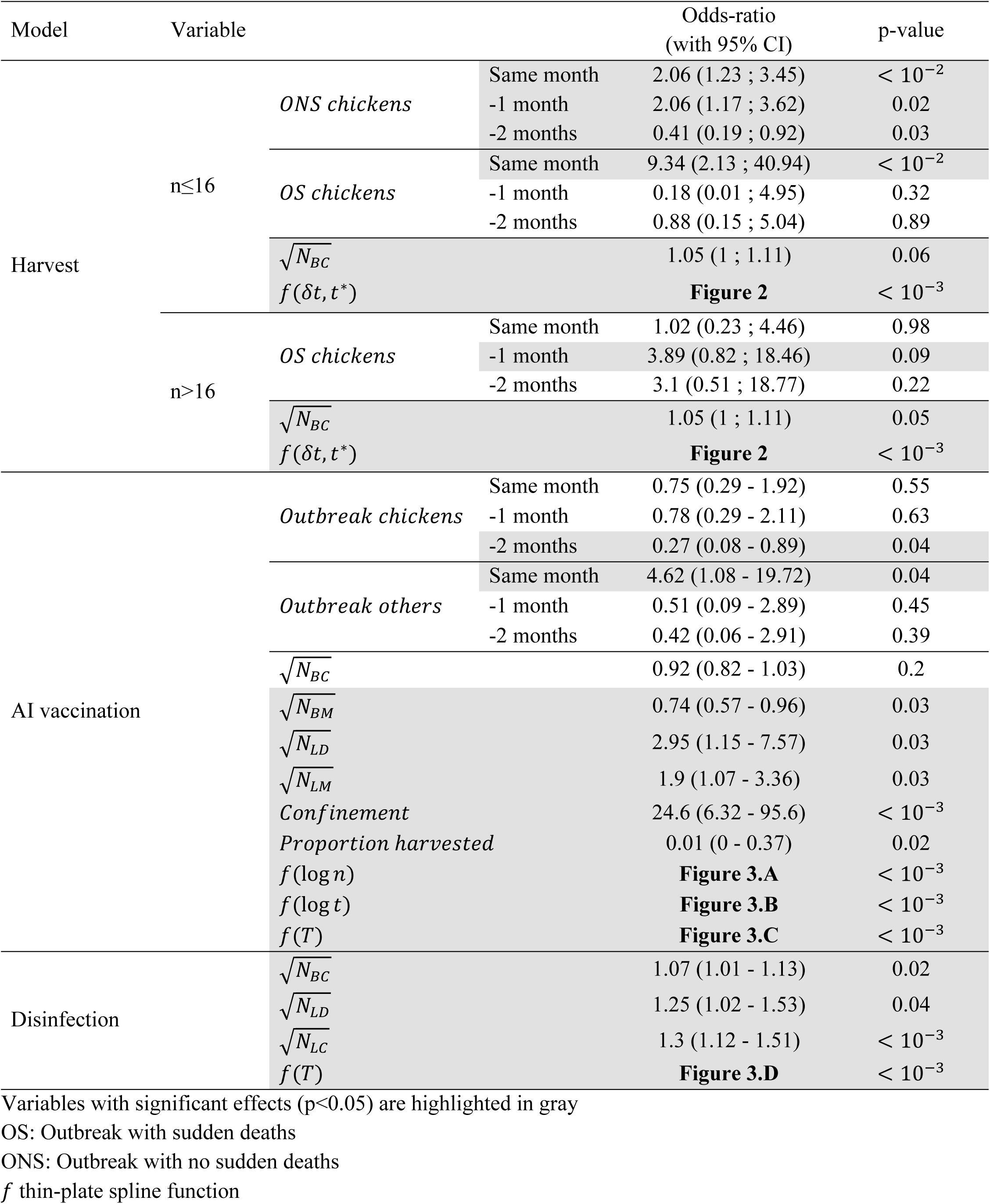
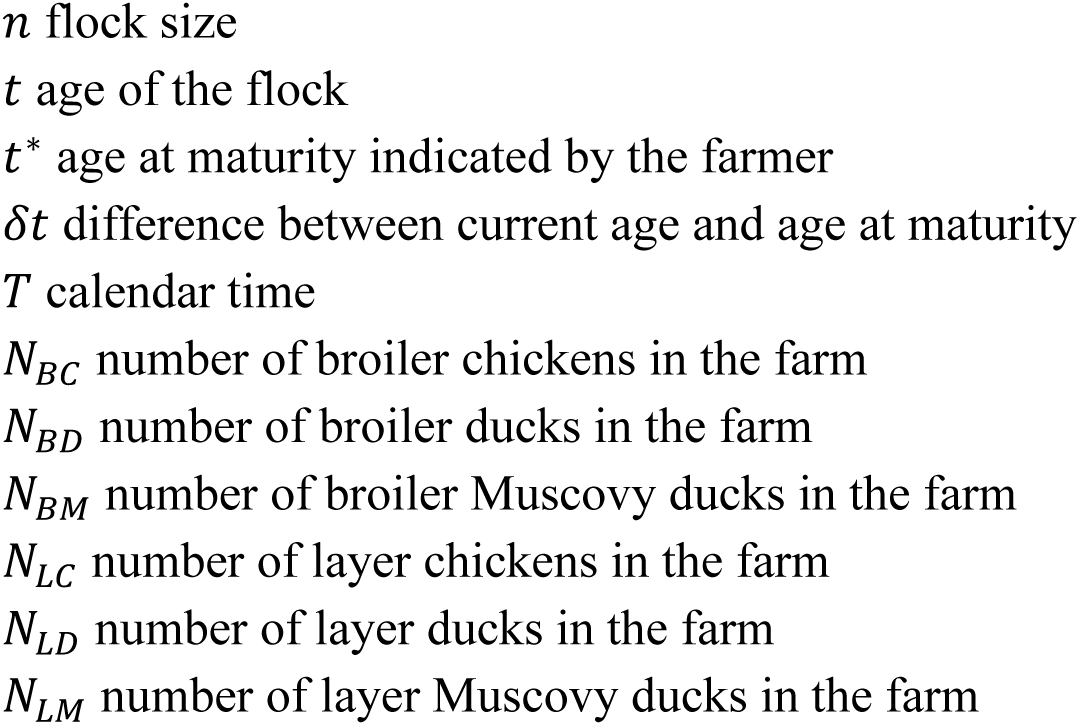
Fitted parameters of the broiler chicken sales model.

**Figure 2.**
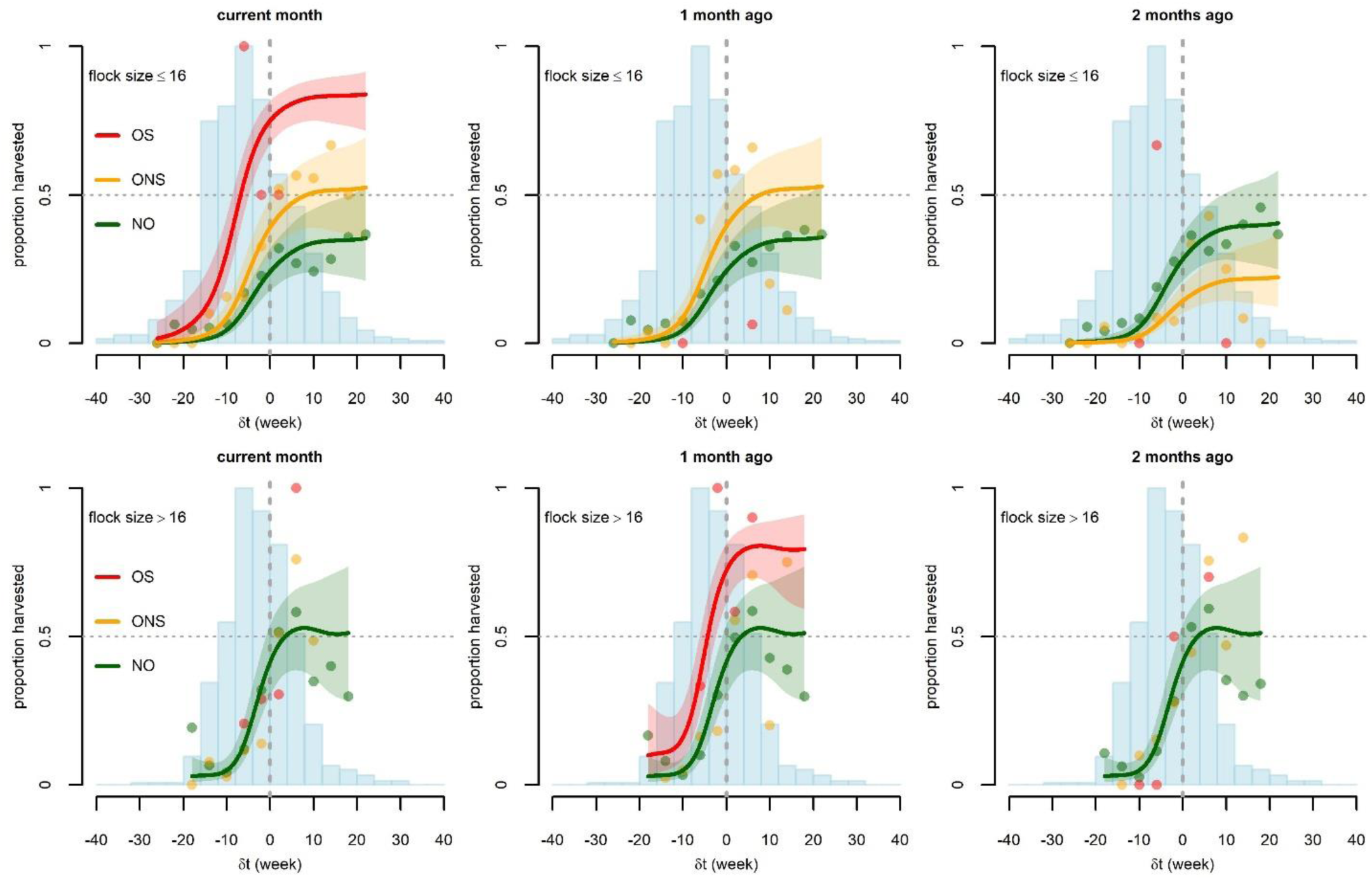
Graphical representation of the relationship between the difference *δt* (current flock age - flock age at maturity) and the proportion of broiler flocks sold in the absence (green color - NO) or presence of outbreaks with disease-induced mortality, either with sudden deaths (red color - OS) or with no sudden deaths (orange color - ONS). Three different outbreak timings are considered: same month (left), one month prior (middle), and two months prior (right). Two different classes of flock size are considered: small (bottom) and large (top). Points are the observed proportions (estimated from at least two flock-months) and lines are the predictions of the fitted Harvest model, along with 90% confidence bands. Model predictions with outbreaks are only displayed when fitted outbreak effects have some statistical significance (*p<0.10*) (see **Table 2**). Blue histograms correspond to the number of observed flock-months in the different classes of *δt* (scaled to their maximum, 139 in the top graphs and 157 in the bottom graphs).

**Figure 3.**
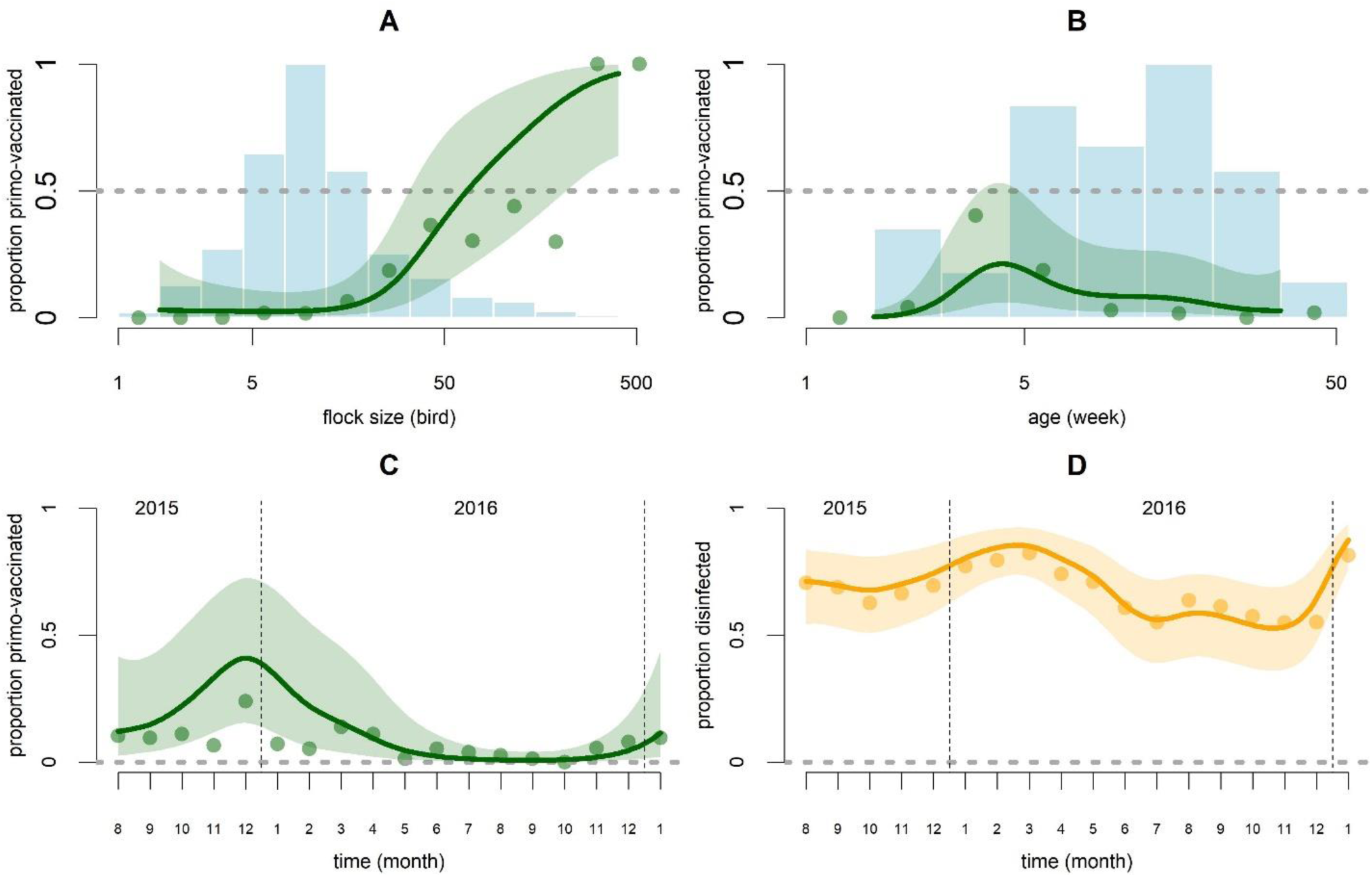
Graphical representation of predictions of the AI vaccination and disinfection models as functions of covariates whose effect is modeled with thin plate smooth splines. For the AI vaccination model (green) these covariates are flock size (n) (A), age (t) (B) and calendar time (T) (C). For the disinfection model (orange), the covariate is calendar time (T) (D). Points are the observed proportions and lines are the predictions along with the 90% confidence band. In graphs C and D the proportions are displayed on the logit scale. Blue histograms correspond to the number of observed flock-months in the different classes of *log (n)* (A) and *t* (B) (scaled to their maximum, 402 in A and 345 in B).

The harvest model explained 34.2% of the observed deviance. Probability of harvest of chicken broiler flocks of the same farm was not shown to be auto-correlated in time As the interaction term between flock size (*n*) and outbreak occurrence is significant (p < 0.01) but difficult to interpret (displayed in **Table S1**), we separated the flocks into large and small. A threshold value of 16 birds per flock gave the lowest AIC (when using a qualitative variable indicating small flock or large flock), and flocks of 16 birds or fewer (52% of all flocks) are referred to as small while flocks of 17 or more (48% of all flocks) are large. As expected, the probability of harvest was found to be strongly dependent on the difference between the flock age and the anticipated age at maturity (*δt*), with older flocks being more likely to be sold. The probability of harvest was close to zero when *δt* < −15 weeks, i.e. flocks that are more than 15 weeks away from being designated as mature. The probability of harvest increased steeply from *δt* = −10 to *δt* = 0. For *δt* > 0 (flocks past their age at maturity), the probability of harvest was consistently high but lower than 100% and did not depend on age. Larger flocks had a steeper increase in harvest probability as a function of *δt*; once past the mature age (*δt* > 0), the estimated probability of harvest for large flocks was higher (41% – 55%) than for small flocks (25% – 41%) (**Figure 2**).

The probability of harvest of small flocks was significantly higher on farms that had experienced a “non-sudden outbreak” (ONS) in chickens in the same month (Odds ratio (OR) = 2.06; 95% confidence interval (CI): 1.23 - 3.45) or the previous month (OR=2.06; 95% CI: 1.17 - 3.62) and was lower in farms that had experienced an ONS in chickens two months prior (OR=0.41; 95% CI: 0.19 - 0.92). The probability of harvest of small flocks was much higher in farms that had experienced a “sudden outbreak” OS in the same month (OR=9.34: 95% CI: 2.13 - 40.94). We used the fitted model to predict the mean harvest proportion in the study population with and without outbreak. Estimated mean harvest proportions of small flocks were 17% (no outbreak), 28% (ONS), and 56% (OS) when considering outbreaks occurring in the same month; this corresponded to harvest increases of 56% and 214% for ONS and OS outbreaks, respectively. Estimated mean harvest proportion was 18% (no outbreak) and 28% (ONS) when considering outbreaks one month prior; this corresponded to a 56% increase in harvest in case of ONS one month prior. Mean harvest proportions were 20% (no outbreak) and 11% (ONS) when considering outbreaks two months prior, indicating a 47% decrease in harvest in case ONS two months prior. For large flocks, ONS in chickens (in any month current or previous) did not have any effect on the harvest of broiler chickens (the removal of ONS variables decreased the model AIC). The occurrence of OS in chickens one month prior may be positively associated with early harvest with an estimated 76% increase in harvest proportion (OR=3.89; 95% CI: 0.82 - 18.46; p=0.09). However, the small sample size – only seven observations of outbreaks with sudden deaths in large flocks – means that we do not have sufficient statistical power to support this association. In the last six months of data collection, farmers were asked to indicate the destination of harvested birds. Based on these partial observations, flocks harvested during or one month after outbreaks in chickens (OS or ONS) were more likely to be sold to traders and less likely to be slaughtered at home (**Table 3**). The likelihood of harvest was also positively correlated with the number of other broiler chickens present on the farm (**Table S1**, p < 0.01). It was not found to be affected by the concomitant introduction of other flocks, vaccination status, or calendar time (T). The farm random effect was significant for large flocks (*σ* = 0.74; 95% CI: 0.47 - 1.17) and not significant for small flocks

**Table 3.**
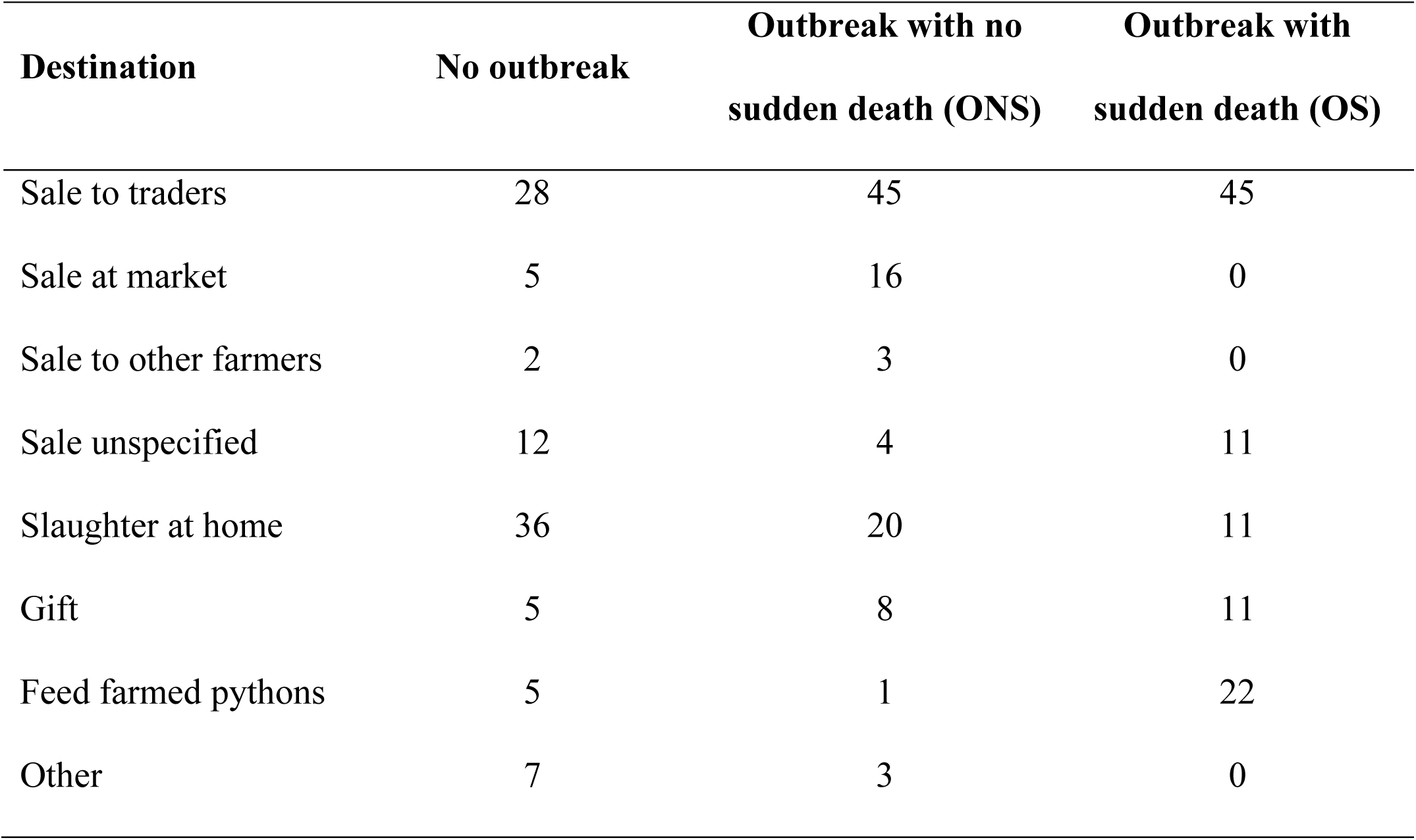
The destination of harvested broiler chicken flocks with or without occurrence of outbreaks of disease-induced mortality in chickens of the same farm in the same month or one month prior (%)

| **Destination** | **No outbreak** | **Outbreak with no sudden death (ONS)** | **Outbreak with sudden death (OS)** |
| --- | --- | --- | --- |
| Sale to traders | 28 | 45 | 45 |
| Sale at market | 5 | 16 | 0 |
| Sale to other farmers | 2 | 3 | 0 |
| Sale unspecified | 12 | 4 | 11 |
| Slaughter at home | 36 | 20 | 11 |
| Gift | 5 | 8 | 11 |
| Feed farmed pythons | 5 | 1 | 22 |
| Other | 7 | 3 | 0 |

In the AI vaccination model, 71.9% of the observations’ deviance was explained. The probability for broiler chicken flocks on the same farm to be vaccinated against AI was not found to be auto-correlated in time. The likelihood of broiler chicken vaccination against AI strongly increased with flock size; probability of vaccination was almost zero for flocks of 16 birds or fewer and nearly 100% for flocks of more than 200 birds (**Figure 3.A**). Vaccination was preferentially performed at 4.3 weeks of age (**Figure 3.B**). Flocks kept indoors or in enclosures had a substantially higher chance of being vaccinated than flocks scavenging outdoors (OR = 24.6; CI: 6.32 - 95.6). Harvested flocks were less likely to receive an AI vaccination (OR = 0.01; CI: 0 - 0.37). The likelihood of AI vaccination varied over time: it increased over the September-January period and decreased during the rest of the year (**Figure 3.C**). The farm random effect was significant (σ = 2.86; CI: 1.88 – 4.35). We failed to obtain convergence when fitting the specific effects of OS and ONS in chickens, so we used an aggregate variable “outbreak in chickens” instead (**Table 2**). Broiler chicken flocks were more likely to be vaccinated if an outbreak had occurred in the same month in other species (OR = 4.62; CI: 1.08 - 19.72; p=0.04) and less likely to be vaccinated if an outbreak had occurred two months prior in chickens (OR = 0.27; CI: 0.08 - 0.89; p=0.03). These two effects were weakly significant and should be interpreted with caution (**Table 2**). The coefficients for interaction terms between outbreak occurrence and flock size were not significantly different from zero. The number of broiler Muscovy ducks present in the farm had a negative effect (p = 0.03) and the number of layer ducks and layer Muscovy ducks had a positive effect (both p = 0.03) on the probability of AI vaccination (**Table 2**)

In the disinfection model, 61.9% of the observations’ deviance was explained. Probability of disinfection on farms was auto-correlated in time (likelihood ratio test for 1-month AR-model on residuals; p < 0.0001); this was not observed for the harvest or vaccination models (both p > 0.3). Consequently, the disinfection model was improved by fitting an AR-1 autoregressive model using the “gamm” routine of the “mgcv” R package (see **technical appendix 4**). The estimated AR-1 autoregressive coefficient was high (*ρ* = 0.71). The likelihood of disinfection of farm facilities increased with the number of layer-breeder hens (OR = 1.3; CI: 1.12 - 1.51; p = 0.001), layer-breeder ducks (OR = 1.25; CI: 1.02 - 1.53; p = 0.03), and to a lesser extent broiler chickens (OR=1.07; CI: 1.01 - 1.13; p = 0.02) present on the farm (**Table 2**). Farm disinfection appeared to have a seasonal component. It was least likely in October-November and most likely in the January-April period (**Figure 3D**). It was not found to be affected by the occurrence of outbreaks on farms (no decrease in AIC when including outbreak occurrence).

## Discussion

Regions like the Mekong river delta combine high human population density, wildlife biodiversity, and agricultural development. As such, they are considered hotspots for the emergence and spread of novel pathogens (*36*). The high density of livestock farmed in semi-commercial operations with limited disease prevention practices further increases the risk of spread of emerging pathogens in livestock and their transmission to humans (*16*). In-depth studies of poultry farmers’ behavioral responses to disease occurrence in animals are needed to understand how emerging pathogens – especially avian influenza viruses – may spread and establish in livestock populations and how optimal management policies should be designed. To the best of our knowledge, this study is the first to provide a detailed and quantified account of the dynamics of livestock management in small-scale farms and its evolution in response to changing epidemiological risks shortly after disease outbreaks occur. While our analysis was performed on a geographically restricted area, the decision-making context of the studied sample of farmers applies to a wide range of poultry producers in low- and middle-income countries. Small-scale poultry farming, combining low investments in infrastructure, no vertical integration, and subject to limited state control on poultry production and trade, is common in most regions affected by avian influenza, in Southeast Asia, Egypt, and West Africa (*37-40*).

In our cohort study, owners of small chicken broiler flocks resorted to early harvesting of poultry, also referred to as depopulation, as a way to mitigate losses from infectious diseases. The revenue earned from the depopulation of flocks might be low, either because birds are still immature or because traders use disease symptoms as an argument to decrease the sale price. Nevertheless, depopulation allows the farmer to avoid a large revenue loss resulting from disease-induced mortality or the costs of management of sick or dead birds. More importantly, farmers avoid the cost of feeding chickens at high risk of dying and prevent the potential infection of subsequently introduced birds. Our results also suggest that the depopulation period, which lasts approximately 2 months, is followed by a “repopulation” period during which farmers lower their harvest rate, possibly to increase their pool of breeding animals in order to repopulate their farm.

The epidemiological effect of chicken depopulation is likely twofold: on the one hand it may slow the transmission of the disease on the farm, since the number of susceptible and infected animals is temporarily decreased; on the other hand, since most poultry harvested during or just after outbreaks were sold to itinerant traders or in markets, depopulation increases the risk of dissemination of the pathogens through trade circuits (*41*). There is epidemiological evidence that poultry farms can be contaminated with HPAI through contact with traders who purchase infectious birds and that infectious birds can contaminate other birds at traders’ storage places and in live bird markets (*12, 14, 42*). Overall, chicken depopulation may reduce local transmission at the expense of long-distance dissemination of the pathogen. The rapid sale of sick birds also exposes consumers and actors of the transformation and distribution chain (traders, slaughterers, retailers) to an increased risk of infection with zoonotic diseases transmitted by poultry, like avian influenza (*43*). Large flocks appear to be less readily harvested upon observation of disease mortality. Farmers may depopulate large flocks only upon observation of sudden deaths, and the number of observation in our study is too small to demonstrate statistical significance of this effect. The likely reason for this difference is that the sale and replacement of larger flocks incurs a higher transaction cost. While small flocks are easily collected and replaced by traders and chick suppliers in regular contact with farmers, the rapid sale of larger flocks probably require the intervention of large-scale traders or several small-scale traders with whom farmers have no direct connection, and who may offer a lower price per bird. When farm production increases, farmers tend to rely on pre-established agreements with traders, middlemen, or hatcheries on the sale dates in order to reduce these transaction costs, giving them little possibility to harvest birds at an earlier time (*44*).

While government-supported vaccination programs have been proposed as a suitable tool to control AI in small scale farms with little infrastructure (*45*), in this cohort AI vaccination was almost exclusively performed in large flocks kept indoors or in an enclosure. Vaccination against AI is believed to be inexpensive for farmers as vaccines are supplied for free by the sub-department of animal health of Ca Mau province and performed by local animal health workers. However, vaccination may still involve some fixed transaction cost as farmers have to declare their flocks to the governmental veterinary services beforehand. Also it is possible that small flocks, being less likely to be sold to distant larger cities (*46*), are less likely to have their vaccination status controlled, making their vaccination less worthwhile from the farmers’ perspective. Crucially, it is these smaller flocks that are more likely to be sold into trading network during outbreaks. Finally, farmers’ willingness to expand their production, invest in farm infrastructure, and implement AI prevention are likely correlated. Farms with a large breeding-laying activity tend to invest more in preventive actions (disinfection and vaccination) compared to farms specialized in broiler production. This may reflect a higher individual market value of layer-breeder hens compared to broiler chicks, making their protection more beneficial.

While vaccination against AI and disinfection appear to depend on individual farmer attitude, as shown by the significance of the farm random effects, they still vary over time when viewed across all farms (**Figure 1**). Contrary to harvesting behavior, these preventive actions have a seasonal component (**Figure 3.C and 3.D**) indicating a willingness to maximize the number of vaccinated broiler chickens and the protection against other diseases during the January-March period. This would be consistent with epidemiological observations as AI transmission increases in the January-February period (*26, 47*). Farm disinfection has a significant temporal autocorrelation component and is unaffected by disease outbreaks, indicating that farmers are slower at adapting this practice to changing conditions.

The last 23 years of emerging pathogen outbreaks and zoonotic transmissions failed to prepare us for the epidemiological catastrophe that we are witnessing in 2020. Multiple subtypes of avian influenza viruses have crossed over into human populations since 1997 (*3, 11*), all resulting from poultry farming activities. Small-scale poultry farming is likely to be maintained in low- and middle-income countries as it provides low-cost protein, supplemental income to rural households, and is supported by consumer preference of local indigenous breeds of poultry (*37, 48, 49*). If we ignore the active role that poultry farmers play in the control and dissemination of avian influenza, we may miss another opportunity to curtail an emerging disease outbreak at a stage when it is still controllable.

## Abbreviations

AIC: Akaike Information Criterion
AI: avian influenza
CI: confidence interval
HPAI: highly pathogenic avian influenza
MGAM: mixed-effects general additive model
ONS: outbreak with no sudden death
OS: outbreak with sudden death
OR: odds ratio

## Author contributions

MFB, NTLT, HTAX, PNT and HML developed the data collection protocol and supervised the data collection; BNVY and AD processed the data; AD developed the main hypotheses, performed the data analysis and wrote the manuscript; MFB, HLM and BNVY reviewed the manuscript; MFB supervised the project and provided a technical support to the data analysis.

## Data availability

The study dataset is available online at https://osf.io/ws3vu/. DOI: 10.17605/OSF.IO/WS3VU

## Supplementary materials

Supplementary materials and methods 1. Selection of observations

Supplementary materials and methods 2. control covariates

Supplementary materials and methods 3. Variable transformation

Supplementary materials and methods 4. Accounting for auto-correlation

Table S1. Original fitted harvest model

## References

1. J. R. Rohr, C. B. Barrett, D. J. Civitello, M. E. Craft, B. Delius, G. A. DeLeo, P. J. Hudson, N. Jouanard, K. H. Nguyen, R. S. Ostfeld, J. V. Remais, G. Riveau, S. H. Sokolow, D. Tilman, Emerging human infectious diseases and the links to global food production. Nat Sustain 2, 445–456 (2019).

2. Y. Guan, J. S. Peiris, A. S. Lipatov, T. M. Ellis, K. C. Dyrting, S. Krauss, L. J. Zhang, R. G. Webster, K. F. Shortridge, Emergence of multiple genotypes of H5N1 avian influenza viruses in Hong Kong SAR. Proc Natl Acad Sci U S A 99, 8950–8955 (2002).

3. R. Gao, B. Cao, Y. Hu, Z. Feng, D. Wang, W. Hu, J. Chen, Z. Jie, H. Qiu, K. Xu, X. Xu, H. Lu, W. Zhu, Z. Gao, N. Xiang, Y. Shen, Z. He, Y. Gu, Z. Zhang, Y. Yang, X. Zhao, L. Zhou, X. Li, S. Zou, Y. Zhang, X. Li, L. Yang, J. Guo, J. Dong, Q. Li, L. Dong, Y. Zhu, T. Bai, S. Wang, P. Hao, W. Yang, Y. Zhang, J. Han, H. Yu, D. Li, G. F. Gao, G. Wu, Y. Wang, Z. Yuan, Y. Shu, Human infection with a novel avian-origin influenza A (H7N9) virus. N Engl J Med 368, 1888–1897 (2013).

4. S. Funk, M. Salathe, V. A. Jansen, Modelling the influence of human behaviour on the spread of infectious diseases: a review. J R Soc Interface 7, 1247–1256 (2010).

5. P. Chilonda, G. Van Huylenbroeck, A conceptual framework for the economic analysis of factors influencing decision-making of small-scale farmers in animal health management. Revue scientifique et technique / Office international des épizooties 20, 687–700 (2001).

6. FAOSTAT. Database. (2019) [cited 2020 Feb 20]; Available from: http://www.fao.org

7. Y. Guan, G. J. D. Smith, The emergence and diversification of panzootic H5N1 influenza viruses. Virus Res 178, 35–43 (2013).

8. OIE. Chapter 10.4. Infection with Avian Influenza viruses. (2018) [cited 2020 Feb 20]; Available from: https://www.oie.int/fileadmin/Home/eng/Health_standards/tahc/current/chapitre_avian_influenza_viruses.pdf

9. M. Imai, T. Watanabe, M. Hatta, S. C. Das, M. Ozawa, K. Shinya, G. Zhong, A. Hanson, H. Katsura, S. Watanabe, C. Li, E. Kawakami, S. Yamada, M. Kiso, Y. Suzuki, E. A. Maher, G. Neumann, Y. Kawaoka, Experimental adaptation of an influenza H5 HA confers respiratory droplet transmission to a reassortant H5 HA/H1N1 virus in ferrets. Nature 486, 420–428 (2012).

10. X. Li, Z. Zhang, A. Yu, S. Y. W. Ho, M. J. Carr, W. Zheng, Y. Zhang, C. Zhu, F. Lei, W. Shi, Global and Local Persistence of Influenza A(H5N1) Virus. Emerg Infect Dis 20, 1287–1295 (2014).

11. S. Lai, Y. Qin, B. J. Cowling, X. Ren, N. A. Wardrop, M. Gilbert, T. K. Tsang, P. Wu, L. Feng, H. Jiang, Z. Peng, J. Zheng, Q. Liao, S. Li, P. W. Horby, J. J. Farrar, G. F. Gao, A. J. Tatem, H. Yu, Global epidemiology of avian influenza A H5N1 virus infection in humans, 1997–2015: a systematic review of individual case data. Lancet Infect Dis 16, e108–e118 (2016).

12. P. K. Biswas, J. P. Christensen, S. S. U. Ahmed, A. Das, M. H. Rahman, H. Barua, M. Giasuddin, A. S. M. A. Hannan, M. A. Habib, N. C. Debnath, Risk for Infection with Highly Pathogenic Avian Influenza Virus (H5N1) in Backyard Chickens, Bangladesh. Emerg Infect Dis 15, 1931–1936 (2009).

13. S. Desvaux, V. Grosbois, T. T. H. Pham, S. Fenwick, S. Tollis, N. H. Pham, A. Tran, F. Roger, Risk Factors of Highly Pathogenic Avian Influenza H5N1 Occurrence at the Village and Farm Levels in the Red River Delta Region in Vietnam. Transbound Emerg Dis 58, 492–502 (2011).

14. N. Y. Kung, R. S. Morris, N. R. Perkins, L. D. Sims, T. M. Ellis, L. Bissett, M. Chow, K. F. Shortridge, Y. Guan, M. J. Peiris, Risk for infection with highly pathogenic influenza A virus (H5N1) in chickens, Hong Kong, 2002. Emerg Infect Dis 13, 412–418 (2007).

15. F. O. Fasina, A. L. Rivas, S. P. R. Bisschop, A. J. Stegeman, J. A. Hernandez, Identification of risk factors associated with highly pathogenic avian influenza H5N1 virus infection in poultry farms, in Nigeria during the epidemic of 2006-2007. Prev Vet Med 98, 204–208 (2011).

16. K. A. Henning, J. Henning, J. Morton, N. T. Long, N. T. Ha, J. Meers, Farm- and flock-level risk factors associated with Highly Pathogenic Avian Influenza outbreaks on small holder duck and chicken farms in the Mekong Delta of Viet Nam. Prev Vet Med 91, 179–188 (2009).

17. B. Hickler, Bridging the gap between HPAI “awareness” and practice in Cambodia. Recommendations from an Anthropological Participatory Assessment. (Food and Agriculture Organisation, Rome, 2007).

18. S. Padmawati, M. Nichter, Community response to avian flu in Central Java, Indonesia. Anthropology & Medicine 15, 31–51 (2008).

19. A. Delabouglise, N. Antoine-Moussiaux, T. D. Phan, D. C. Dao, T. T. Nguyen, B. D. Truong, X. N. Nguyen, T. D. Vu, K. V. Nguyen, H. T. Le, G. Salem, M. Peyre, The Perceived Value of Passive Animal Health Surveillance: The Case of Highly Pathogenic Avian Influenza in Vietnam. Zoonoses Public Health 63, 112–128 (2016).

20. R. Sultana, N. Ali Rimi, S. Azad, M. S. Islam, M. S. Uddin Khan, E. S. Gurley, N. Nahar, S. P. Luby, Bangladeshi backyard poultry raisers’ perceptions and practices related to zoonotic transmission of avian influenza. J Infect Dev Ctries 6, 156–165 (2012).

21. L. Zhang, T. Pan, Surviving the crisis: Adaptive wisdom, coping mechanisms and local responses to avian influenza threats in Haining, China. Anthropology & Medicine 15, 19–30 (2008).

22. A. Hidano, G. Enticott, R. M. Christley, M. C. Gates, Modeling Dynamic Human Behavioral Changes in Animal Disease Models: Challenges and Opportunities for Addressing Bias. Frontiers in Veterinary Science 5, (2018).

23. D. R.-H. a. D. Z. Jennifer Ifft, Production and Risk Prevention Response of Free Range Chicken Producers in Viet Nam toHighly Pathogenic Avian Influenza Outbreaks. Am J Agr Econ 93, 490–497 (2011).

24. A. Hidano, M. C. Gates, Why sold, not culled? Analysing farm and animal characteristics associated with livestock selling practices. Prev Vet Med 166, 65–77 (2019).

25. A. Delabouglise, B. Nguyen-Van-Yen, N. T. L. Thanh, H. T. A. Xuyen, P. N. Tuyet, H. M. Lam, M. F. Boni, Poultry population dynamics and mortality risks in smallholder farms of the Mekong river delta region. Bmc Vet Res 15, (2019).

26. A. Delabouglise, M. Choisy, T. D. Phan, N. Antoine-Moussiaux, M. Peyre, T. D. Vu, D. U. Pfeiffer, G. Fournié, Economic factors influencing zoonotic disease dynamics: demand for poultry meat and seasonal transmission of avian influenza in Vietnam. Sci Rep 7, (2017).

27. General Statistics Office of Vietnam, Results of the 2016 Rural, Agriculture and Fishery Census. (Vietnam Statistical publishing House, Hanoi, Vietnam, 2017).

28. OIE. World Animal Health Information Database (WAHIS Interface). (2019) [cited 2020 Feb 20]; Available from: http://www.oie.int/wahis_2/public/wahid.php/Wahidhome/Home

29. N. T. L. Thanh, N. H. T. Vy, H. T. A. Xuyen, H. T. Phuong, P. N. Tuyet, N. T. Huy, B. Nguyen-Van-Yen, H. M. Lam, M. F. Boni, No Evidence of On-farm Circulation of Avian Influenza H5 Subtype in Ca Mau Province, Southern Vietnam, March 2016 - January 2017. PLoS Curr 9, (2017).

30. J. C. Mariner, B. A. Jones, S. Hendrickx, I. El Masry, Y. Jobre, C. C. Jost, Experiences in participatory surveillance and community-based reporting systems for H5N1 highly pathogenic avian influenza: a case study approach. Ecohealth 11, 22–35 (2014).

31. S. N. Wood, N. Pya, B. Säfken, Smoothing Parameter and Model Selection for General Smooth Models. Journal of the American Statistical Association 111, 1548–1563 (2017).

32. S. Wood, Generalized Additive Models: an Introduction with R. (Chapman and Hall/CRC, Boca Raton, Florida, ed. 2, 2017).

33. B. Naimi, N. A. S. Hamm, T. A. Groen, A. K. Skidmore, A. G. Toxopeus, Where is positional uncertainty a problem for species distribution modelling? Ecography 37, 191–203 (2014).

34. D. W. Hosmer, S. Lemeshow, Applied Logistic Regression. W. A. Shewart, S. S. Wilks, Eds., (John Wiley & Sons, Hoboken, New Jersey, 2000).

35. R core team. R: a language and environment for statistical computing. R Foundation for Statistical Computing (2014) [cited 2012 Oct 8]; Available from: http://www.R-project.org/

36. T. Allen, K. A. Murray, C. Zambrana-Torrelio, S. S. Morse, C. Rondinini, M. Di Marco, N. Breit, K. J. Olival, P. Daszak, Global hotspots and correlates of emerging zoonotic diseases. Nat Commun 8, 1124 (2017).

37. A. Sudarman, K. M. Rich, T. Randolph, F. Unger, Poultry value chains and HPAI in Indonesia: The case of Bogor. Africa/Indonesia Team Working Paper No. 27 (Food and Agriculture Organisation of the United Nations, Rome, 2010).

38. T. U. Obi, A. Olubukola, G. A. Maina, Pro-poor HPAI risk reduction strategies in Nigeria Africa/Indonesia Team Working Paper No. 5 (Food and Agriculture Organisation of the United Nations, Rome, 2008).

39. F. A. Hosny, The structure and importance of the commercial and village based poultry systems in Egypt. (Food and Agriculture Organisation of the United Nations, Rome, 2006).

40. S. Burgos, J. Hinrichs, J. Otte, D. Pfeiffer, D. Roland-Holst, S. K. T. O., Poultry, HPAI and Livelihoods in Cambodia - A Review., Mekong Team Working Paper No. 3 (Food and Agriculture Organisation of the United Nations, Rome, 2008).

41. A. Delabouglise, M. F. Boni, Game theory of vaccination and depopulation for managing livestock diseases and zoonoses on small-scale farms. Epidemics, 100370 (2019).

42. G. Fournié, A. Tripodi, T. T. T. Nguyen, V. T. Nguyen, T. T. Tran, A. Bisson, D. U. Pfeiffer, S. H. Newman, Investigating poultry trade patterns to guide avian influenza surveillance and control: a case study in Vietnam. Sci Rep 6, 29463 (2016).

43. G. Fournié, E. Hoeg, T. Barnett, D. U. Pfeiffer, P. Mangtani, A Systematic Review and Meta-Analysis of Practices Exposing Humans to Avian Influenza Viruses, Their Prevalence, and Rationale. Am J Trop Med Hyg 97, 376–388 (2017).

44. M. A. O. Catelo, A. C. Costales, Contract Farming and Other Market Institutions as Mechanisms for Integrating Smallholder Livestock Producers in the Growth and Development of the Livestock Sector in Developing Countries. PPLPI Working Paper No. 45 (Food and Agriculture Organisation of the United Nations, Rome, 2008).

45. FAO, Approaches to controlling, preventing and eliminating H5N1 Highly Pathogenic Avian Influenza in endemic countries. (Food and Agriculture Organization of the United Nations, Rome, 2011).

46. D. X. Tung, A. Costales, Market Participation of Smallholder Poultry Producers in Northern Viet Nam., (Food and Agriculture Organisation of the United Nations, Rome, 2007).

47. L. O. Durand, P. Glew, D. Gross, M. Kasper, S. Trock, I. K. Kim, J. S. Bresee, R. Donis, T. M. Uyeki, M. A. Widdowson, E. Azziz-Baumgartner, Timing of influenza A(H5N1) in poultry and humans and seasonal influenza activity worldwide, 2004-2013. Emerg. Infect. Dis. 21, 202–208 (2015).

48. S. Burgos, S. Hinrichs, J. Otte, D. Pfeiffer, D. Roland-Holst, Poultry, HPAI and Livelihoods in Viet Nam - A Review., Mekong Team Working Paper No. 2 (Food and Agriculture Organisation of the United Nations, Rome, 2008).

49. M. Epprecht, Geographic Dimensions of Livestock Holdings in Vietnam. Spatial Relationships among Poverty, Infrastructure and the Environment. PPLPI Working Paper No. 24 (Ministry of Agriculture and Rural Development of Vietnam, Hanoi, Vietnam, 2005).

