## Supplementary materials and methods for "Poultry farmer response to disease outbreaks in smallholder farming systems"

**Supplementary materials and methods 1. selection of observations**

We included three distinct binary variables for the occurrence of an outbreak in the same month, one month prior, and two months prior. For this reason, observations made in the two first months of the study were discarded since, during these two months, it was unknown whether farms had previously experienced outbreaks.

In the “disinfection” model, observations were farm-months. We removed farm-month with missing data on disinfection performed by farmers (18 farm-months). 858 farm-months were used to fit the disinfection model. In the “harvest” and “AI vaccination”, observations were chicken broiler flock-months. We selected all chicken flock-months more than 10 days old at the time of data collection and classified by farmers as "broilers". A total of 1656 flock-months were available for inclusion in the model. Additional specific selection was performed for the harvest and vaccination models.

1. Harvest model

We removed flock-months which less than 20 days old at the time of data collection. This 20-day threshold was chosen because some newborn flocks below this age were partly sold, not for meat consumption but for management on other farms. Also, we removed flock-months where no chickens were available for harvest because they had all died in the course of the month (25 flock-months). In total, 153 flock-months were removed and 1503 flock-months were used to fit the harvest model.

2. AI vaccination model

We removed flock-months of flocks which had already been vaccinated against avian influenza in a previous month, since vaccination is usually performed only once (among the 338 vaccinated flocks, only 8 were vaccinated a second time). We also removed flock-months whose housing conditions were not reported (4 flock-months). In total, 338 flock-months were removed and 1318 flock-months were used to fit the AI vaccination model.

**Supplementary materials and methods 2. complete models and control covariates**

For the harvest model, the main control variable is, logically, (1) the body weight of chickens, as broiler chickens are conventionally harvested after a fattening period upon reaching a given weight. Since the chicken weight was not collected during the survey, we used the difference between the current flock age $t$ and the anticipated age at maturity $t^*$ indicated by farmers in the questionnaire. Hereafter we use $\delta t = t - t^*$ for this difference. The shape of the function linking $\delta t$ and harvest may depart from linearity and is affected by the chicken breed, which determines the growth performance. Since information on chicken breed was not collected we used the age at maturity $t^*$ and the flock size ($n$) as proxy indicators of the growing performance of the breed and built a proxy body weight variable as a multivariate spline function of $\delta t$, $t^*$ and $n$ (*1*). 20% of flock-months had missing value for t*. Since there was little within-farm variation in t* (2 months of difference at most between two flocks of the same farm), missing values were replaced by the median t* in the other flocks of the corresponding farm. (2) The calendar time $T$ was included as an additional smoothing spline term, since harvest may also be influenced by market prices which vary from one month to the other. Control variables included as standard linear terms were (1) the number of chickens kept for laying eggs or breeding ($N_{LC}$) - famers with a large breeder-layer activity may want to keep some broilers chickens in the farm for replacing the breeding-laying stock, making them less likely to harvest broilers; (2) the number of broiler chickens simultaneously present in the same farm in other flocks ($N_{BC}$); (3) the number of chicken flocks introduced in the same month; (4) the number of chicken flocks introduced in the previous month – farmers with a high number of broilers chickens or many recently introduced broiler flocks may want to sell their current flocks faster in order to limit feeding expenses and workload; (5) the vaccination status of the flock against AI; (6) the vaccination status of the flock against Newcastle Disease (ND) – farmers may keep their vaccinated flocks for a longer period as they are at lower risk of being affected by an infectious disease. we assumed the effect of outbreaks on the dependent variable may be affected by the size of the considered flock (n). Consequently, we included an interaction term between outbreaks and *log(n)* in the analysis.

For the AI primo-vaccination model, control variables included as smoothing splines were (1) the flock age *t* - vaccination may be preferentially done early in the flock life, and (2) the flock size *n* and (3) the calendar time *T* - vaccination activities may be intensified at particular times of the year. Control variables included as standard linear terms were (1-6) the size of populations of broiler and layer-breeder chickens, ducks and Muscovy ducks - farmers' perceived risk of AI and attitude towards vaccination may be influenced by the size of the poultry population at risk for AI; (7) the type of housing (free-range or confinement in pens or indoor) which affects the convenience of vaccination; and (8) the proportion of the flock harvested in the same month - farmers might be less willing to vaccinate flocks being harvested. we assumed the effect of outbreaks on the dependent variable may be affected by the size of the considered flock (n). Consequently, we included an interaction term between outbreaks and log(n) in the analysis.

For the disinfection model, control variables included as smoothing splines were (1) the calendar time T - disinfection activities may be intensified at particular times of the year. Control variables included as standard linear terms were (1-6) the size of populations of broiler and layer-breeder chickens, ducks and Muscovy ducks - the farmers' attitude towards prevention may be influenced by the size of the poultry population at risk of disease.

**Supplementary materials and methods 3. variable transformation**

Since flock age *t*, flock size *n*, and farm poultry population size variables were highly skewed (see **Table 1** in the main text), we transformed them in different ways for inclusion in the model.

*t* and *n* being strictly positive, they were log-transformed. Farm populations of broiler and layer-breeders of different species being null or positive, they were square-root transformed. Covariates included in the multivariate spline function for body weight ($\delta t$, $t^*$, log $n$) were centered and standardized. In the case of the harvest model, since the effect of outbreaks was highly influenced by the flock size, in order to simplify the visualization and interpretation of the results we fitted an additional piecewise MGAM model on two sub-population: small and large flocks. The threshold value for flock size used to differentiate the two sub-populations was determined by testing all possible thresholds and selecting the one returning the lowest AIC (Akaike Information Criterion).

**Supplementary materials and methods 4. accounting for auto-correlation**

Arguably, one farmer is likely to maintain the same farm management from one month to the next despite changes in influential covariates. Therefore, for each model, we tested the presence of farm-level time autocorrelation by fitting two linear regression models on the deviance residuals, with a fixed constant effect and with and without intra-farm AR-1 time autocorrelation structure and comparing the two model fits with a log-likelihood ratio test. If the fit was significantly improved by including the autocorrelation term, we implemented the same model fitting protocol with an additional intra-farm AR-1 time autocorrelation term on the dependent variable. We used the "gamm" routine of the "mgcv" package for this purpose (*2*). Since "gamm" models for binomial data are fitted with the penalized quasi-likelihood approach, the AIC metric is not suitable to compare such models. Instead, we implemented a stepwise removal of covariates whose t-test returned the highest probability of type 1 error (p-value) until all remaining covariates had a p-value lower than 20%.

**References**


1. S. Burgos, S. Hinrichs, J. Otte, D. Pfeiffer, D. Roland-Holst, *Poultry, HPAI and Livelihoods in Viet Nam - A Review.*, Mekong Team Working Paper No. 2 (Food and Agriculture Organisation of the United Nations, Rome, 2008).
2. S. Wood, *Generalized Additive Models: an Introduction with R*. (Chapman and Hall/CRC, Boca Raton, Florida, ed. 2, 2017).
