## Supplementary material for "Poultry farmer response to disease outbreaks in smallholder farming systems": Table S1

**Table S1. Original fitted harvest model**

| Model | Variable | | Odds-ratio (with 95% CI) | p-value |
|---|---|---|---|---|
| Harvest | $ONS\ chickens$ | Same month | 1.39 (0.95 ; 2.04) | 0.09 |
| | | -1 month | 1.53 (1.03 ; 2.27) | 0.04 |
| | | -2 months | 0.69 (0.43 ; 1.09) | 0.11 |
| | $ONS\ chickens * \log\ n$ | Same month | 0.54 (0.37 ; 0.79) | $< 10^{-2}$ |
| | | -1 month | 0.75 (0.51 ; 1.1) | 0.14 |
| | | -2 months | 1.57 (1.03 ; 2.41) | 0.04 |
| | $OS\ chickens$ | Same month | 3.73 (1.36 ; 10.26) | 0.02 |
| | | -1 month | 1.95 (0.58 ; 6.49) | 0.28 |
| | | -2 months | 1.49 (0.42 ; 5.24) | 0.54 |
| | $OS\ chickens * \log\ n$ | Same month | 0.37 (0.14 ; 0.99) | 0.05 |
| | | -1 month | 0.97 (0.25 ; 3.74) | 0.97 |
| | | -2 months | 1.11 (0.28 ; 4.36) | 0.89 |
| | $\sqrt{N_{BC}}$ | | 1.07 (1.03 ; 1.11) | $< 10^{-2}$ |
| | $f(\delta t, t^*, \log\ n)$ | | | $< 10^{-3}$ |

$f$ thin-plate spline function

$n$ flock size

$t^*$ age at maturity indicated by the farmer

$\delta t$ difference between current age and age at maturity

$N_{BC}$ number of broiler chickens in the farm
